# Alteration of skin fibroblast steady state contributes to healing outcomes

**DOI:** 10.1101/2024.12.06.627278

**Authors:** Yaqing Huang, Nuoya Wang, Hao Xing, Jingru Tian, Dingyao Zhang, Daqian Gao, Henry C. Hsia, Jun Lu, Micha Sam Brickman Raredon, Themis R. Kyriakides

## Abstract

Fibroblasts display complex functions associated with distinct gene expression profiles that influence matrix production and cell communications and the autonomy of tissue development and repair. Thrombospondin-2 (TSP-2), produced by fibroblasts, is a potent angiogenesis inhibitor and negatively associated with tissue repair. Single-cell (sc) sequencing analysis on WT and TSP2KO skin fibroblasts demonstrate distinct cell heterogeneity. Specifically, we found an enrichment of Sox10+ multipotent progenitor cells, identified as Schwann precursor cells, in TSP2KO fibroblasts, while fibrosis-related subpopulations decreased. Immunostaining of tissue and cells validated the increase of this Sox10+ population in KO fibroblasts. Furthermore, in silico analysis suggested enhanced pro-survival signaling, including WNT, TGF-β, and PDGF-β, alongside a reduced BMP4 response. Additionally, the creation of two TSP2KO NIH3T3 cell lines using the CRISPR/Cas9 technique allowed functional and signaling validation in a less complex system. Moreover, KO 3T3 cells exhibited enhanced migration and proliferation, with elevated levels of pro-regenerative molecules including TGF-β3 and Wnt4, and enrichment of nuclear β-catenin. These functional and molecular alterations likely contribute to improved healing and increased neurogenesis in TSP2-deficient wounds. Overall, our findings describe the heterogeneity of dermal fibroblasts and identify pro-regenerative features of TSP2KO fibroblasts.

## Introduction

Skin wounds in mammals heal by fibrosis, a repair mechanism that involves the rapid deposition of collagen and other extracellular matrix components that contribute to scar formation [1]. This fibroproliferative response, while effective in preventing infection and fluid loss, often results in scar tissue formation that differs structurally and functionally from the original skin. This is due to excess deposition of collagen fibrils that leads to thickening of skin with altered mechanical properties, and contributes to its inability to regenerate complex tissue structures like hair follicles, sweat glands. It is appreciated that multiple cell types, including inflammatory cells and fibroblasts, play critical roles in determining healing outcomes.

Fibroblasts are major extracellular matrix (ECM) producers and perform functions critical to wound healing [2]. Upon injury, activated fibroblasts migrate to the wound site and proliferate during the early phase of healing. Subsequently, they differentiate into myofibroblasts that contract the wound and synthesize new ECM, thus facilitating tissue repair. Fibroblasts also secrete various signaling molecules that recruit other cell types, such as macrophages, epithelial cells, and endothelial cells, to coordinate the wound healing process. Control of diverse fibroblast functions, including proliferation, differentiation, ECM production, and their communication with other cells, can influence healing outcomes.

In addition to functional diversity, fibroblasts demonstrate molecular heterogeneity across and within organs, which is also critical for tissue homeostasis [2–5]. For example, subsets of skin and intestine fibroblasts contribute to tissue renewal via remodeling of the ECM and serving as a stem cell niche for hair follicles or intestinal crypts, respectively [6, 7]. These subpopulations are characterized by the expression of WNT signaling components that support epithelial and hair follicle stem cell maintenance [6–9]. In contrast, fibroblasts in tissues with lower cellular turnover, such as the lung and kidney, exhibit fewer stem-like markers but are nonetheless vital for repair, primarily by secreting signaling molecules [2]. It is appreciated that tissue-specific heterogeneity is associated with distinct functions in homoeostasis and following injury. Within tissues, fibroblast subpopulations can be categorized based on their spatial distribution. For example, cells near the upper layer of skin are defined as papillary fibroblasts (Pdgfra+CD26+Sca-), whereas cells in the reticular dermis are called reticular fibroblasts (Pdgfra+Dlk1+Sca-) [10]. Papillary and reticular fibroblasts produce distinct extracellular matrices consisting of non-fibrillary collagens and proteoglycans and well-organized fibrillary collagen bundles, respectively [11, 12]. Due to the capability to deposit collagenous ECM, reticular fibroblasts are considered more fibrotic whereas papillary fibroblasts are more pro-<u>regenerative</u> [13]. Thus, the balance between these two populations can determine the outcome of skin wound healing [10]. Additionally, fibroblasts with specific molecular signatures can exhibit distinct behaviors in tissue repair. For example, En1+ and Cthrc1+ fibroblasts are associated with fibrosis [14, 15], while Wnt1+ and Lef1+ fibroblasts are linked to reduced scarring [15, 16].Therefore, <u>elucidating fibroblast identities and behaviors is essential for understanding the lack of regeneration in issue repair processes.</u>

Thrombospondin-2 (TSP2) is an ECM matricellular glycoprotein, whose levels of expression are negatively correlated with tissue repair outcomes. For example, mice with increased or low/null TSP2 levels display delayed or accelerated wound healing, respectively. Primarily, TSP2 exerts anti-angiogenetic effects via binding to CD36 and CD47 receptors on endothelial cells [17, 18]. Additionally, TSP2 has been implicated in ECM assembly as it was shown that TSP2 deficiency disrupts collagen fibril morphology and organization [19]. Moreoever, TSP2 deficient wounds display increased levels of soluble growth factors and enzymes, such as vascular endothelial growth factor (VEGF) and matrix metalloproteinases (MMPs) [20]. Collectively, changes in ECM assembly and remodeling, along with changes in angiogenesis and growth factor release, suggest that TSP2 plays a pivotal role in wound healing. Indeed, TSP2KO wounds display accelerated epithelization, increased angiogenesis, and reduced scar formation [20, 21]. Wounds treated with anti-TSP2 constructs also exhibit improved healing with enhanced cell infiltration and angiogenesis, and the emergence of new nerve bundles [22–25]. Collectively, these findings underscore a strong connection between TSP2 and cellular aspects of generation. Differences between the healing capacities of WT and TSP2 KO skin, coupled with our understanding of fibroblasts as the major source of TSP2, prompted us to evaluate these cells in an unbiased high throughput manner. Specifically, we applied single-cell (sc-) and bulk RNA sequencing on WT and TSP2KO mouse dermal fibroblasts. In silico analyses indicated an increased Sox10+ population in KO fibroblasts, which was validated by immunostaining of isolated cells and tissue. Moreover, ECM-related categories and cell functions, including proliferation and migration, were identified as the top regulated pathways in KO fibroblasts. A potential crosstalk between WNT, BMP, and TGF-β signaling was suggested, which may contribute to the more regenerative phenotype of TSP2KO wounds such as enhanced neurogenesis. Additionally, we employed CRISPR/Cas9 technology to precisely eliminate TSP2 expression and study the subsequent effects in a less heterogeneous cell population. Functional assays demonstrated enhanced proliferation and migration of TSP2KO fibroblasts. Molecular analyses revealed increased levels of WNT4 and TGF-β3 and enrichment in nuclear β-catenin in KO cells. Collectively, our findings highlight the heterogeneity of skin fibroblasts and their programming towards a fibrotic response. Importantly, investigation of TSP2-deficient cells reveals a shift of the cell steady state in TSP2KO skin towards pro-regenerative signaling.

## Method and Materials

### Animals

C57bl/6J and TSP2KO mice aged 12 to 14 weeks were used. Only male mice were used for sequencing. C57bl/6J mice were purchased from Jackson Laboratory. TSP2KO mice were generated as described previously 18. All animal study procedures were approved by the Yale Institutional Animal Care and Use Committee (IACUC). All mice were kept in a 12-hour light/dark environment and fed ordinary feed. All authors complied with ARRIVE guidelines.

### Dermal Fibroblasts Isolation and Sequencing Sample Preparation

Dermal fibroblasts (DFs) were isolated from 12-week-old WT and TSP2KO mouse dorsal skin according to the established protocol. Briefly, mouse dorsal skins were shaved, excised, treated with 5% antibiotic-antimycotic (Gibco), and incubated in 25 μg/ml Trypsin (Sigma) overnight at 4°C. After separation from the epidermis and adipose tissue, the dermis was physically disrupted and then digested using collagenase IV (Worthington Biochemical) for 3 hrs under 37 °C. After centrifuge, cell pellet was then resuspended in 25mM glucose Dulbecco’s Modified Eagle Medium (DMEM, Gibco) with 10% (v/v) fetal bovine serum (FBS, Peak Serum Inc) and 1% (v/v) penicillin-streptomycin (P/S, Gibco) and culture for 3-5 days to enrich the fibroblast population.

For single-cell RNA sample preparation, after the enrichment assay, fibroblasts were lifted using trypsin (Sigma), suspended in PBS at a concentration of 1000 cells/µl, and passed through a 100 µm cell strainer (BD Bioscience). The single-cell suspension was sent to the Yale Microarray Center, where library preparation was performed using the Chromium Next GEM single-cell 3’ v3 reagent kit (10x Genomics). The libraries were then sequenced on the Illumina NovaSeq 6000 platform with a 150 bp read length.

For bulk RNA sequencing, RNA was isolated from passage 1 fibroblasts using the RNeasy Mini Kit (Qiagen) according to the manufacturer’s protocol. The RNA concentration was determined using a Nanodrop spectrophotometer (Thermo Scientific). Approximately 2 µg of RNA per sample was sent to the Yale Genomics Core for sequencing using the Illumina TruSeq Stranded mRNA kit (Illumina).

### Sequencing Data Analysis

Sequencing data were aligned to the mouse genome (mm10) using Cell Ranger (10x Genomics). The raw matrix was then analyzed with the Seurat R package (version 5.0.3) [26]. Cells with low-quality reads (nFeatures < 300, or mitochondrial gene percentage < 0.1% or > 10%) were excluded during the QC process. A total of 18,193 and 20,134 cells remained for WT and KO samples (n=3), respectively, and these were merged and integrated using canonical correlation analysis (CCA) reduction (dims.use = 1:40). Graph-based clustering was performed using the FindClusters function in Seurat (resolution = 0.4, dims.use = 1:40), and markers for each cluster were identified using the FindAllMarkers function (min.pct = 0.5, log fold change > 0.25). Cluster visualization on a 2D map was done using uniform manifold approximation and projection (UMAP) via the RunUMAP function [27].

Given the uneven distribution of cell numbers across clusters, differential gene analysis between genotypes within individual clusters was performed to minimize confounding variance. Specifically, comparisons were made between individual KO and WT samples, resulting in the generation of nine gene lists. A scoring value was created to reflect the frequency of each gene across the nine gene lists. P-values were integrated using Fisher’s method. Genes were ranked based on Fisher p-value, average fold change (Avg Log2FC), ratio, power (Avg Log2FC × ratio), and score (frequency across datasets). Genes with a score of 6 or higher were selected for further analysis. Downstream GO pathway and network analysis were performed using the STRING database (https://string-db.org/).

For cell-cell communication analysis, cells were processed using the NICHES package [28]. Autocrine signaling in endothelial, epithelial, and immune cells were excluded to focus specifically on fibroblast interactions. The signaling archetypes were annotated based on the sending and receiving cell types, with most interactions being related to fibroblast autocrine pathways. Cell-cell signaling edges were down-sampled to mitigate sample variation due to sample size and integrated with rpca (resolution = 0.2, dims.use = 29). Differential cell-cell signaling between genotypes was explored using the FindMarkers function.

Violin, ridge, and feature plots of specific gene expression levels were generated using the corresponding functions in Seurat. Other visualizations, such as dot plots, histograms, and heatmaps, were generated using the ggplot2 and pheatmap packages [29, 30].

For bulk RNA-seq data, FASTQ files from individual samples were inputted and analyzed using the Partek Flow platform. Specifically, reads were aligned to the mouse reference genome (mm10) using STAR within the platform [31]. The aligned reads were then mapped to known transcripts using RefSeq as the annotation model. Low-expression genes (features < 10) were filtered out, and the remaining counts were normalized based on the median ratio (DESeq2 only). Following normalization, principal component analysis (PCA) and differential analysis were performed. A list of 462 genes was generated using thresholds of FDR < 0.05 and fold change > 1.5 or < -1.5. Downstream pathway analysis, including GO pathways and GSEA, was conducted using the STRING database (https://string-db.org/) and the Web-based Gene Set Analysis Toolkit (https://www.webgestalt.org).

### Flow Cytometry (FACS)

A total of 1,000,000 isolated fibroblasts were fixed using 2% PFA (J.T. Baker) for 20 minutes and permeabilized with 0.1% Triton-X in PBS for 15 minutes. Cells were treated with 1% BSA in PBS for non-specific blocking and then incubated with FACS antibodies. Untreated fibroblasts were used as a negative control, and beads stained with corresponding antibodies were used as a positive control. All cells were passed through a 100 µm cell strainer (BD Bioscience) before being loaded onto the FACSAria II (BD Bioscience), and signals were recorded for 10^5 events. Laser parameters were set based on the negative and positive controls. Data analysis was then performed using FlowJo software (FlowJo, LLC).

### Immunofluorescence (IF) Staining

For wound tissue IF staining, 4-mm thick paraffin-embedded tissue slides from WT and TSP2KO mouse D7 wounds were processed according to standard protocols. Primary antibody was diluted in 1% BSA in PBS and applied to the tissue area. After overnight incubation at 4°C, slides were incubated with secondary antibody solution with DAPI for 1 hr at RT and mounted using VECTASHIELD® Antifade Mounting Medium (Vector Laboratories, H-1000-10). Stained tissues were then imaged under EVOS Cell Imaging System (ThermoFisher Scientific). Images were analyzed with ImageJ (Molecular Devices).

### CRISPR TSP2KO Cell lines Creation

CRISPR/Cas9 Knock out experiments were conducted using LentiCRISPR(pXPR_001) plasmid as directed by the protocol [32]. Briefly, four guideRNA targeting TSP2 geno sequences were designed (sequence information was enclosed in **Table S3.1**) and inserted into the vector. To make lentivirus, lentiCRISPR was co-transfected into HEK293T cells with packaging plasmids pVSVg (Addgene 8454) and psPAX2 (Addgene 12260). Next, NIH3T3 cells were infected using the produced lentivirus with or without TSP2 guide RNA. Infected cells were then processed via single cell sorting and plated into individual well in 96-well plate to obtain single cell colony. The depletion of protein was validated using western blot.

### Reverse Transcription and quantitative-RT-PCR

Reverse transcription was completed with the QuantiTect Reverse Transcription Kit (Qiagen). Quantitative real-time PCR (qRT-PCR) was performed with the iTaq Universal SYBR Green One-Step Kit (Bio-Rad) on a CFX96 Touch Real-Time PCR machine (Bio-Rad). Primers were ordered from the Keck Oligonucleotide Facility (Yale University) from sequences found in PrimerBank (Harvard University) or published literature (**Table S1**). Data were normalized to GAPDH or β-actin expression.

### Western Blot

Lysates were extracted from cells using RIPA buffer (pH = 7.4, Boston BioProducts, Inc.) supplemented with cOmplete EDTA-free protease inhibitor cocktail (Roche). Protein concentration was determined using BCA protein assay kit according to supplier’s instructions (Thermofisher). Standard western blot was then performed using a mini-PROTEAN TGX Stain-Free Gel (10%, Bio-Rad) and the Licor Odyssy CLx system was used to visualize blots. Densitometry analysis was performed using ImageJ Gels plugin.

### Immunofluorescence (IF) Staining

For cell IF staining, samples were fixed in 4% paraformaldehyde (PFA, J.T. Baker), permeabilized using 0.1% Triton-X, immersed with 1% BSA in PBS for blocking, and incubated with primary antibody for overnight at 4°C and secondary antibody at RT for 1 hr subsequently. Stained cells were then imaged under EVOS Cell Imaging System (ThermoFisher Scientific). Images were analyzed with ImageJ (Molecular Devices).

### Scratch Assay

Cell scratch assay was used to evaluate the migration ability and conducted based on the protocol described previously ^17^. Briefly, fibroblasts were seeded in a 6-well plate and allowed to grow to 100 % confluence. A scratch was created in each well with a 200 μL pipet tip. The cells were then washed with PBS and incubated in fresh media. The plates were then monitored and photographed using a Zeiss Axio Vert.A1 inverted light microscope for the following 24 hours. The area of the scratch at each time point was measured with ImageJ, and relative wound closure (%) calculated for each well. All experiments were performed at least three times in triplicate.

### Cell Proliferation

Fibroblast proliferation was examined using Cell Counting Kit-8 (CCK-8, ab228554, Abcam) according to the manufacture protocol. Specifically, a total number of 3×10^3^ CRISPR Control and TSP2KO NIH 3T3s were seeded in a 96-well plate per well. Cell proliferation rate was determined in cultures supplemented with 10% CCK-8 using a plate reader (O.D. 460) 10% CCK-8 solution.

### Cell Transfection

Cell transfection was performed using transfectamine 5000 (AAT Bioquest) or lipofectamine 3000 (Thermo Fisher) according to the suppliers’ protocola. Briefly, TSP2KO CRISPR cells were seeded on a 6-well plate the day before transfection at a concentration of 2×10^4^. Once reach 80% confluency, reagents and pcDNA plasmid were thawed in advance, then mixed at the ratio of 2.5µg DNA: 7.5 µL transfectamine 5000, or 2.5µg DNA: 3.75 µL lipofectamine 3000: 5 µL P3000 in Optimen. After sitting in room temperature for 20 mins, the DNA and lipofectamine mixture was added into individual plates with fresh media. After 48-72 hr incubation, media was changed to normal media with 750 µg/ml G418 (Gibco), and cells are cultured for another 48 hrs to select out those without transfection. Then cells were processed either by western blot or immunostaining for molecular validation and further analysis. Cells transfected with pcDNA3.1 GFP were utilized to assess transfection efficiency. For β-catenin stanning, pcDNA3.1 c was used as a control vector instead.

### Reagents

The following primary antibodies were used for flow cytometry, IHC or IF: Sox10 Antibody [Alexa Fluor 700] (1:200, Novus Biologics, NBP2-59621AF700), anti-Neurofilament H (NFH) (1:200, EMD Millipore, AB1989), anti-Vimentin (1:500, EMD Millipore, AB5733), TGFB3 Rabbit pAb (1:250, ABclonal, A8460), Sox10 Rabbit pAb (1:100, ABclonal, A15100), β-catenin (IF: 1:20, R&D system, AF1329).

The following primary antibodies were used for western blot: TSP2 Antibody (1:250, GenScript), β-Catenin Antibody (1:500, Cell Signaling, 9562S), Anti-LOX antibody (1:500, Abcam, ab31238), Human/Mouse Wnt4 antibody (1:500, R&D system, MAB475), TGFB3 Rabbit pAb (1:500, ABclonal, A8460), GAPDH Rabbit mAb (1:1000, Cell Signaling, 5174S), and anti-β-actin (1:1000, Abcam, ab8226), HSP90 Anbody (1:1000, Santa Cruz Biotechnology, sc-13119)

The following secondary antibodies were used for immunofluorescence staining or flow cytometry: anti-rabbit IgG Alexa Fluor 488 (1:1000, Invitrogen, a11008), anti-chicken IgG Alexa Fluor 488 (1:1000, Abcam, ab150169), anti-rabbit IgG Alexa Fluor 555 (1:1000, Abcam, ab150078), anti-goat IgG Alexa Fluor 647 (1:500, Invitrogen, a21447), and anti-goat IgG Alexa Fluor 488 (1:1000, Invitrogen, a21467). The following secondary antibodies were used for western blots in the Licor Odyssy DLx system: anti-mouse IgG Alexa Fluor 680 (1:1000, Invitrogen, a10038), and anti-rabbit IgG Alexa Fluor 800 (1:1000, Invitrogen, a32735).

### Statistical Analyses

Error bars represent the standard deviation (SD) unless stated otherwise. An unpaired Student’s t-test (two-tailed) was performed for comparisons between two groups. For comparisons involving three or more groups, one-way analysis of variance (ANOVA) with Tukey’s post hoc test for multiple comparisons was used. Statistical methods other than these two were specified with the results. All statistical analyses were performed using GraphPad Prism 9 or in R. Non-significant comparisons are not shown in the figures by default.

## Results

### Single-cell RNA transcriptomics reveals distinct WT and TSP2KO dermal fibroblast heterogeneity

Isolated dermal fibroblasts (DFs) from WT and TSP2KO mouse back skin (male, 12-week-old, n=3) were subjected to single-cell RNA sequencing (scRNA-seq) (**Figure 1A**). After assessing sample quality based on UMI counts, gene features, and mitochondrial percentage, we obtained 20,134 cells from KO and 18,193 cells from WT samples. Using classical markers such as Col1a1, Ptprc, Epcam, and Cdh5, we identified four major cell types in the skin: mesenchymal, immune, epithelial, and endothelial cells (**Figure S1**). Cell proportion analysis revealed >95% fibroblast abundance across samples, confirming the reliability of the fibroblast enrichment protocol (**Figure S1**).

**Figure 1.**
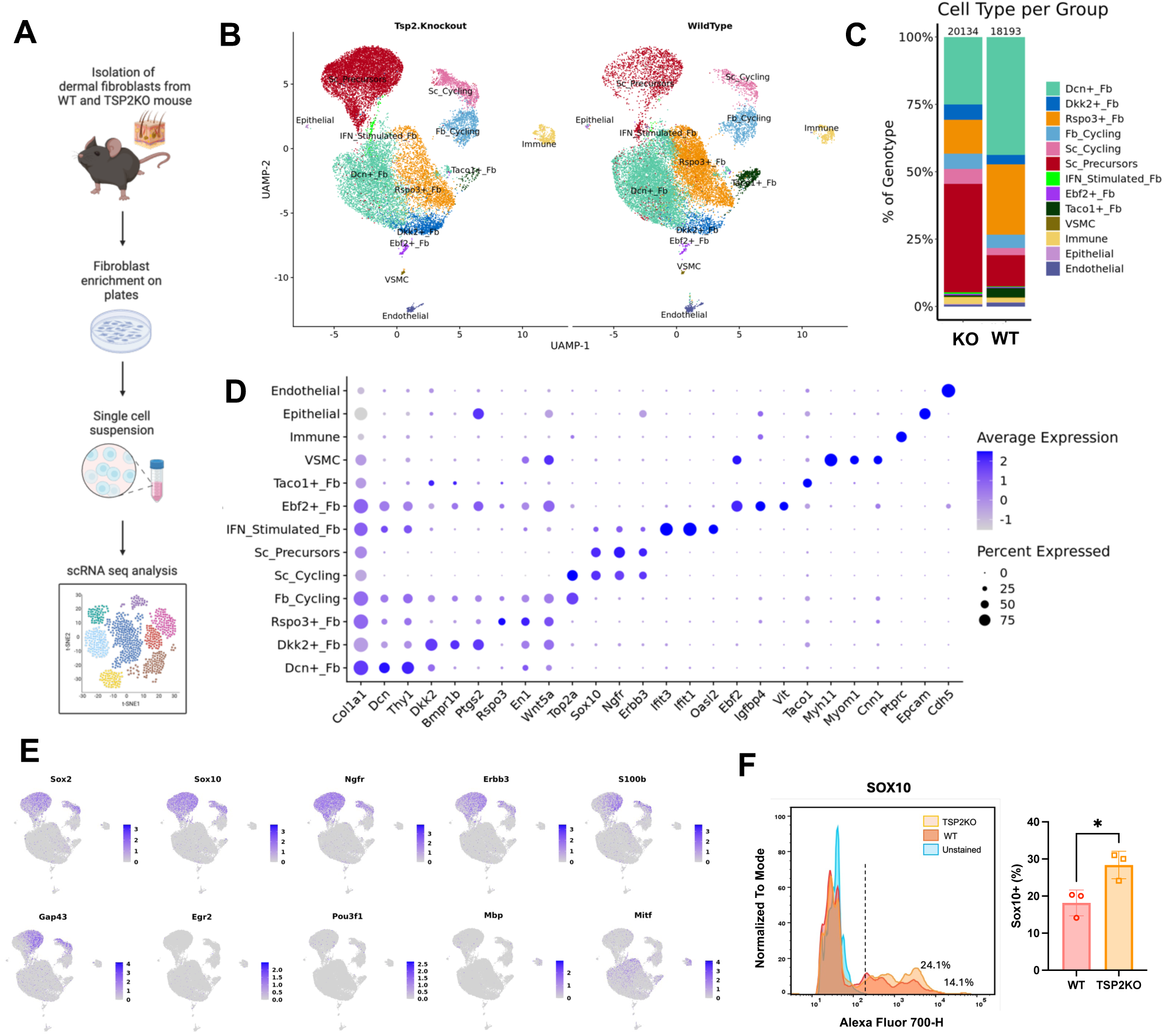
Single cell transcriptomics reveals distinct WT and TSP2KO dermal fibroblast heterogeneity. (A) Diagram of dermal fibroblast isolation and sequencing process; (B) Cell cluster UMAP plot; (B) Cell type distribution in each genotype; (D) Dot plot of selected marker genes for each cell cluster; (E) Feature plots of representative markers for Schwann cell development within Sox10+ population; (F) Histogram of Sox10-700 intensity of unstained and stained dermal fibroblasts and quantification of Sox10+ cells in WT and TSP2KO groups (n = 3, Unpaired T test, two-tailed, *, p<0.05).

Within the Col1a1+ population, we identified 10 distinct cell subtypes based on specific marker expression: DCN+, Rspo3+, Dkk2+, Ebf2+, Taco1+, IFN_Stimulated fibroblasts (Fb), Schwann Precursors (ScP), Vascular smooth muscle cells (VSMC), and Schwann (Sc) or Fibroblast (Fb) Cycling cells (Top2a+) (**Figure 1**).

The main fibroblast populations, Dcn+, Rspo3+, and Dkk2+, decreased in KO group. They express common fibroblast markers such as Thy1 (CD90), Pdgfra, S100a4, Dlk1, and Ly6a (Sca) with ECM-related genes, including Adamts2, Dkk3, Lox, and Fbn1, enriched in the Dcn+ population. Rspo3+ fibroblasts were highly positive for En1, Ltbp1, Wnt5a, and Fgf10, genes associated with WNT and TGF-β signaling pathways. In contrast, Dkk2+ fibroblasts expressed Bmpr1b, Ptgs1, Ptgs2, and Mmp3, constituting a novel signaling profile.

A notable finding was the increased proportion of a Sox10+ population in the KO samples (**Figure 1**), in which various neural crest stem cell makers including Sox2, Sox10, and Ngfr were present. Flow cytometry analysis of mouse skin fibroblasts confirmed this increased population of Sox10+ cells in the TSP2KO group (**Figure 1**). Moreover, this population expressed Schwann-cell-lineage markers like Erbb3, S100b, and Gap43 instead of melanocyte markers like Mitf. However, they were negative for the mature Schwann cell markers like Egr2, Pou3f1, and Mbp (**Figure 1 & S1**). Thus, we believe this population remains in a more dedifferentiated state and named it Schwann Precursors (ScP). Cycling cells were identified using markers such as Top2a and Ccna2.

Unlike major fibroblast populations that displayed overlap in multiple genes, smaller subsets showed patterns in the expression of specific markers. For example, Taco1, which encodes a mitochondrial protein that functions as a translational activator of mitochondrially-encoded cytochrome c oxidase 1, was only expressed in Taco1+ fibroblasts. Interestingly, this population, though small in WT, almost disappeared in KO fibroblasts (**Figure S1**). IFN-Stimulated fibroblasts expressed a combination of interferon (IFN) -stimulated genes including Ifit1, Ifit3, and Oasl2, suggesting a potential role in immune response for acute infection. Ebf2, a highly selective marker for brown and beige fat precursor cells [33], was present in two cell populations. Nevertheless, none of these populations expressed other adipocyte precursor cell markers like Klf4 and Pparg (**Figure S1**). Instead, one was positive for smooth muscle cell markers like Myh11, Myom1, and Cnn1 thus identified as vascular smooth muscle cell (VSMC). The other was then named Ebf2+ fibroblasts with Igfbp4 and Vit enrichment.

Taken together, clustering analysis demonstrates the heterogeneity of skin fibroblasts and suggests a shift in cell state distribution with TSP2 depletion. Specifically, cells expressing standard skin fibroblast markers such as Pdgfra, Thy1, En1, Dcn, Rspo3 are diminished in the KO population. In contrast, a population resembling Schwann precursor cells is increased in the KO group.

### Cell-cluster specific gene differential analysis highlights shifted cell states with the loss of TSP2

To further explore the differences among fibroblast subpopulations, we conducted a cluster-specific differential gene analysis. In all clusters, TSP2 was among the top downregulated genes in KO samples, confirming the dataset’s reliability. Similarly, osteoglycin (Ogn), a small keratan sulfate proteoglycan, was consistently downregulated in all KO clusters except Schwann precursors (**Figure 2 and Table S2-6**). In the largest fibroblast subpopulations (Dcn+ and Rspo3+), homeobox C8 (Hoxc8), a transcription factor essential for growth and differentiation, and growth factor augmenter of liver regeneration (Gfer), a protein critical for mitochondrial biogenesis, were downregulated in KO samples. In contrast, genes such as Grem1, Sod2, Tnc, Cebpb, and Snrfp were commonly upregulated in the Dcn+, Rspo3+, and Dkk2+ populations (**Figure 2 and S2.2**). These upregulated genes in the KO group are involved in crucial biological processes and cellular homeostasis. For instance, Grem1 is a BMP antagonist and VEGFR agonist to regulate cell differentiation and angiogenesis [34, 35]; Sod2 encodes mitochondrial superoxide dismutase 2, which protects cells from oxidative stress; Tnc (tenascin-C), as described before, is a matrix glycoprotein mediating cell and ECM interactions; Cebpb is a transcription factor that is involved in cell proliferation and differentiation; and Snrfp encodes a protein that participates in pre-mRNA splicing and RNA binding. In addition to these shared genes, Dcn+ fibroblasts demonstrated more growth factor- and extracellular matrix-related genes that are highly differential expressed between WT and KO samples including Igfbp2, Fgf10, Col18a1, Postn, Col5a3, Col4a5, Col8a1, Col11a1, and Efemp1. These genes are related to morphogenesis, matrix organization, and cell adhesion per GO pathway enrichment analysis. Rspo3+ cells displayed downregulation of Sfrp1 (secreted frizzled-related protein 1), a WNT inhibitor, indicating a potential negative effect of TSP2 in WNT activation. Interestingly, Trp53, a gene that induces cell cycle arrest and apoptosis is increased in KO, which may contribute to the decreased percentage of Rspo3+ population. Consequently, GO analysis highlighted fibroblast proliferation, locomotion, and migration as top biological processes affected by the loss of TSP2.

**Figure 2.**
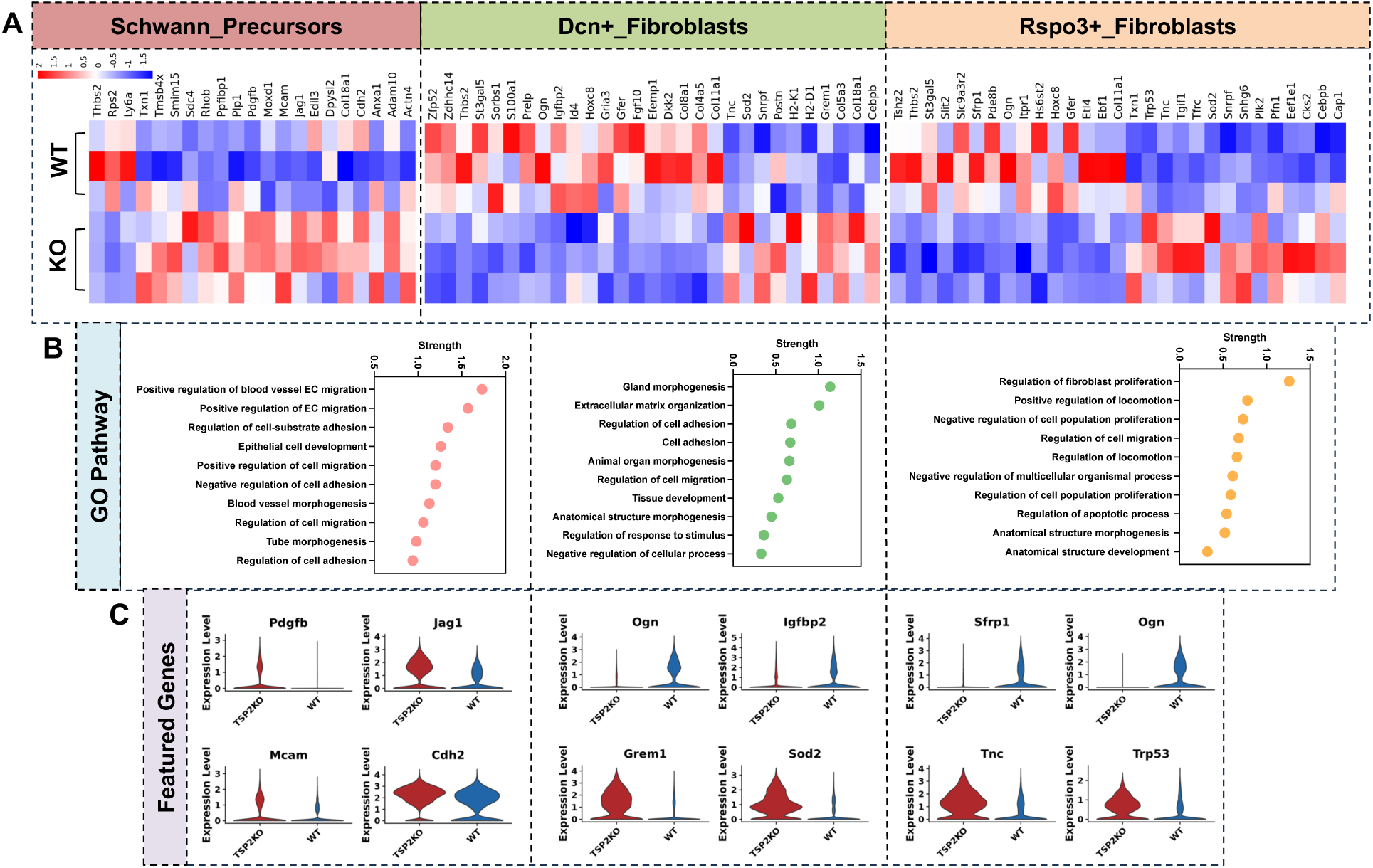
Cell-cluster-specific gene differential analysis identifies highly regulated genes by TSP2 depletion in each cluster. (A) Heatmap of highly differential genes (score ≥ 6, |power| > 1), (B) GO pathway analysis, and (C) violin plots of highly regulated genes between WT and KO within Schwann Precursors, DCN+, and Rspo3+ fibroblasts.

Schwann Precursors in KO group expressed elevated levels of many genes that are pro-survival and pro-angiogenic. The top 10 upregulated genes included Pdgfb (Platelet-derived growth factor subunit), Mcam (Cell surface glycoprotein MUC18), Moxd1 (monooxygenase protein), Jag1 (Jagged-1), Plp1 (Myelin proteolipid protein), Cdh2 (Calcium-dependent cell adhesion protein), Col18a1, Sdc4 (Cell surface proteoglycan), Edil3 (EGF like repeats and discoidin domains 3), Anxa1 (Annexin A1) (**Figure 2 and Table S2**). Most of those play a positive role in cell proliferation, migration, survival, and fate determination via both autocrine and paracrine pathways. Specifically, Pdgfb is known for its effect in promoting cell survival and proliferation. Jag1 is a potent Notch ligand which can activate a series of biological processes including angiogenesis [36]. Mcam and Cdh2 are critical for cell adhesion thus regulating cell interactions. Edil3 is essential in angiogenesis and vascular development and Annexin-A1 is important for VEGF-mediated endothelial cell migration. GO analysis identified that positive regulation of endothelial cell migration was the top regulated pathway, followed by cell-substrate adhesion and epithelial cell development.

Overall, cluster-specific differential analysis emphasizes the heterogeneous effects of TSP2 deficiency on fibroblast function and molecular expression. Nevertheless, common features, including alterations in BMP and WNT signaling related molecules, are observed across major fibroblast populations. Additionally, various cell function-related genes such as Pdgfb, Jag1, and Cdh2 are involved in both autocrine and paracrine interactions, pointing to potential disruptions in cell-cell communication.

### Cell-cell communication analysis indicates enhanced PDGFb and reduced Bmp4 signaling in TSP2KO fibroblasts

To investigate cell-cell signaling, we applied the NICHES method, which analyzes communication between individual pairs of cells (referred to as edges), to our dataset. Signaling involving Schwann Precursors and Schwann Cycling cells was enriched in the KO population (**Figure 3**). Ligand-receptor communications involving Erbb3 and Ngfr were particularly prominent for them as receiving population. Signaling received by fibroblast populations was marked by mechanisms involving FGF receptors, such as Fgf5-Fgfr1, Fgf10-Fgfr1, and Pf4-Fgfr2. Signaling via these pathways originated from Schwann Precursors, other fibroblasts, and immune cells, respectively (**Figure 3**).

**Figure 3.**
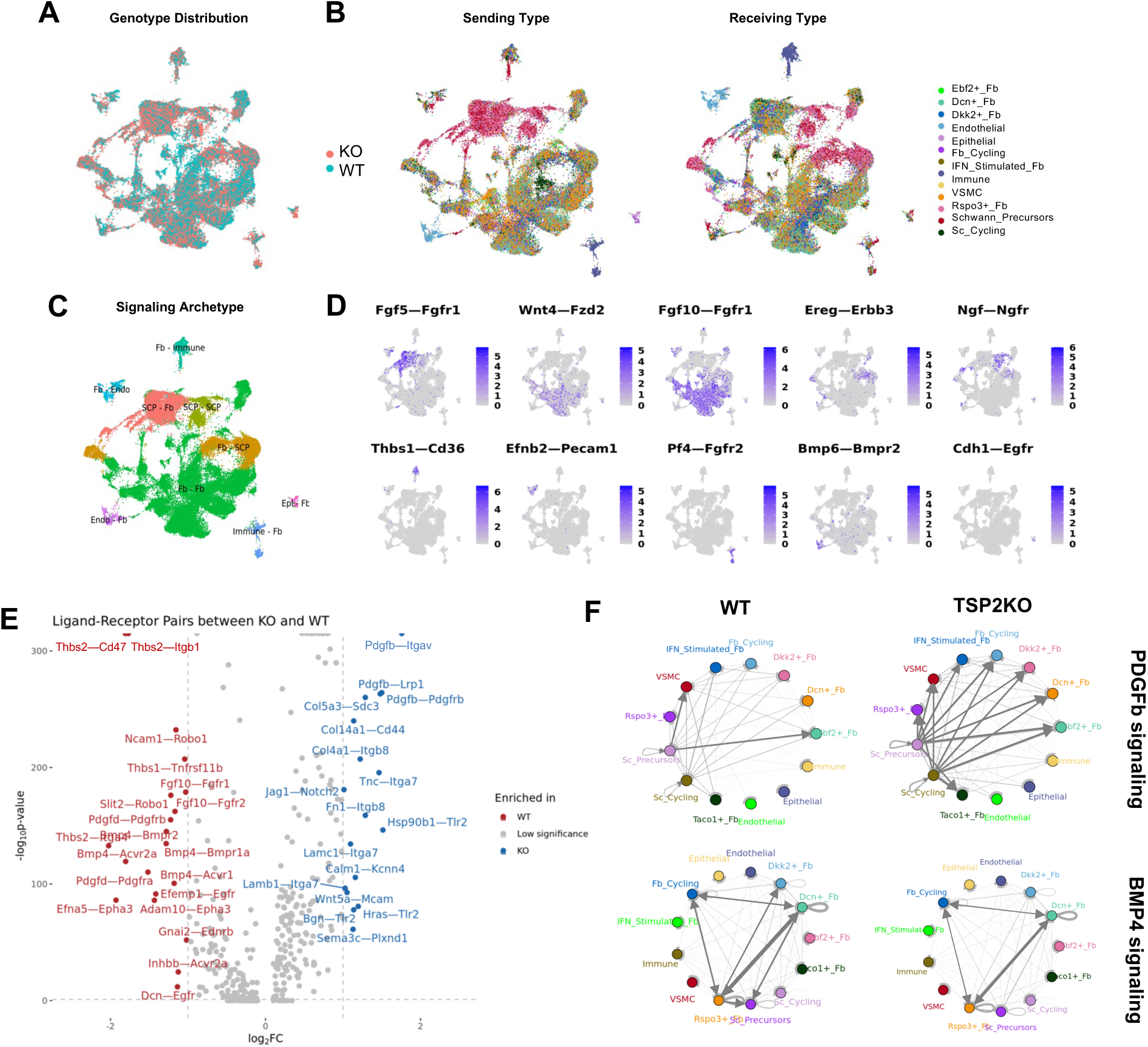
NICHE analysis indicates enhanced PDGFb and reduced Bmp4 intercellular signaling in TSP2KO fibroblasts. (A) The distribution of WT and KO signaling edges in a 2D UMAP plot; (B) Sending cell type and receiving cell type of each edge; (C) Signaling Archetypes defined via Louvain clustering; (D) Representative signaling mechanism for each cluster; (E) Volcano plot of highly regulated ligand-receptor mechanisms perturbed by KO; (F) Circuit plot of PDGFb and Bmp4 family signaling in WT and KO cells (edge thickness is proportional to mean connectivity and frequency, respectively).

Enrichment of communications between WT and KO fibroblasts showed difference due to both perturbed cell numbers and molecule expression. For example, Wnt5a-Mcam (Fb–ScP) and Fgf10-Fgfr1/2 (Fb–Fb), were enriched in either KO or WT populations due to differences in ScP and traditional fibroblast population. TSP2-related signaling was downregulated in KO samples as expected. Notably, Bmp4 signaling pathways, including Bmp4-Bmpr1a/Bmpr2/Acvr2/Acvr1 are enriched in WT group with Dcn+ and Rspo3+ cells being the major producers and receivers. Therefore, the reduction of BMP4 communication in KO might be due to the decreased percentage of major fibroblast population. In contrast, PDGFb-related signaling, including Pdgfb-Lrp1, Pdgfb-Pdgfrb, and Pdgfb-Itgav, and Notch signaling was upregulated in KO fibroblasts (**Figure 3 and S3**). Circuit plot delineating cell-cell communications demonstrated that Schwann Precursors and Schwann Cycling cells were the major senders of PDGFb signaling, with more cells involved and enhanced signaling intensity represented by the thickness of the edges (**Figure 3**).

Overall, cell-cell communication analysis suggests enhanced PDGFb and reduced Bmp4 signaling in TSP2KO cells with altered cell players. These analyses are aligned with previous observations such as increased PDGFb expression in KO Schwann Precursors and enhanced Bmp4 antagonist, Grem1, in Dcn+ fibroblasts. Given the roles of PDGFb and BMP signaling in cell proliferation, migration, and differentiation[37–39], TSP2KO fibroblasts likely exhibit altered behavior in these processes.

### Bulk RNA sequencing confirms perturbation of ECM-related proteins and suggests a positive regulation of WNT pathway in KO fibroblasts

To investigate global transcriptional changes caused by TSP2 deficiency in fibroblasts, we performed bulk-RNA sequencing, which is more sensitive to rare transcripts, on RNA isolated from DFs. We identified 462 significantly perturbed genes and confirmed changes in ECM genes and cell migration. ECM proteins identified by both sequencing including Cspg4, Tnc, Adamts2, Postn, Dcn, Tgfbi, and Lum, are known to contribute to ECM assembly and cell-ECM interactions. Interestingly, DCN knockout ECM exhibits similar structural changes as TSP2KO fibrils [40]. Previous proteomic data also showed decreased levels of Dcn, Lum, and Tgfbi in KO skin ECM, so we aimed to validate their expression levels first. However, qPCR and western blot results were inconsistent for Dcn, Tgfbi, and Lum, likely due to cell heterogeneity. This heterogeneity was also emphasized in the Tissue expression pattern of the global genes where Schwann cell, Microglia, Bone-marrow derived macrophages and glial cells are suggested to be top compartments affected (**Figure S4**).

Intriguingly, Gene Set Enrichment Analysis (GSEA) indicated a positive regulation of cell-cell WNT signaling in addition to vascular-related processes in KO fibroblasts, implicating a new molecular pathway affected by loss of TSP2. Western blot and qPCR on dermal fibroblasts confirmed a robust increased expression of TGF-β3 and WNT4, two molecules that are involved in WNT signaling, in KO group. Furthermore, pathways involving interleukin-1 receptor activity and immune responses were downregulated in KO fibroblasts, suggesting a potential role for TSP2 in inflammation.

In summary, bulk RNA seq analysis confirms perturbation in ECM proteins and cell functions and hints a positive regulation of WNT signaling globally after incorporates cell heterogeneity. These findings are in consistent with established understanding of TSP2 function but also suggest new molecular pathways regulated by TSP2, for example, WNT4 and TGF-β3.

### Depletion of TSP2 in NIH3T3 cells using CRISPR/Cas9 allows functional exploration in a less heterogeneous system

Given the heterogeneity observed in both *in vivo* environments and isolated primary dermal fibroblasts, we used CRISPR/Cas9 to deplete TSP2 in NIH 3T3 cells to create a less complex system. Briefly, we designed four guide RNAs targeting different regions of the TSP2 gene and used virus-encoded delivery to infect NIH 3T3 cells. After single cell sorting, we obtained colonies for two KO cell lines (KO-1 and KO-2) and one control cell line infected with an empty vector (**Figure 5**). Western blot analysis of cell lysates and media from the two TSP2 KO cell lines confirmed successful TSP2 depletion. Additionally, a downstream target of TSP2, Lox, was reduced in KO samples, providing further validation of the knockout (**Figure 5**). qPCR analysis showed no significant difference in Sox10 and Ngfr mRNA levels between KO and control cell lines (**Figure S5**), indicating that TSP2 depletion does not alter cell fate in NIH3T3s, and that the population is homogeneous.

**Figure 4.**
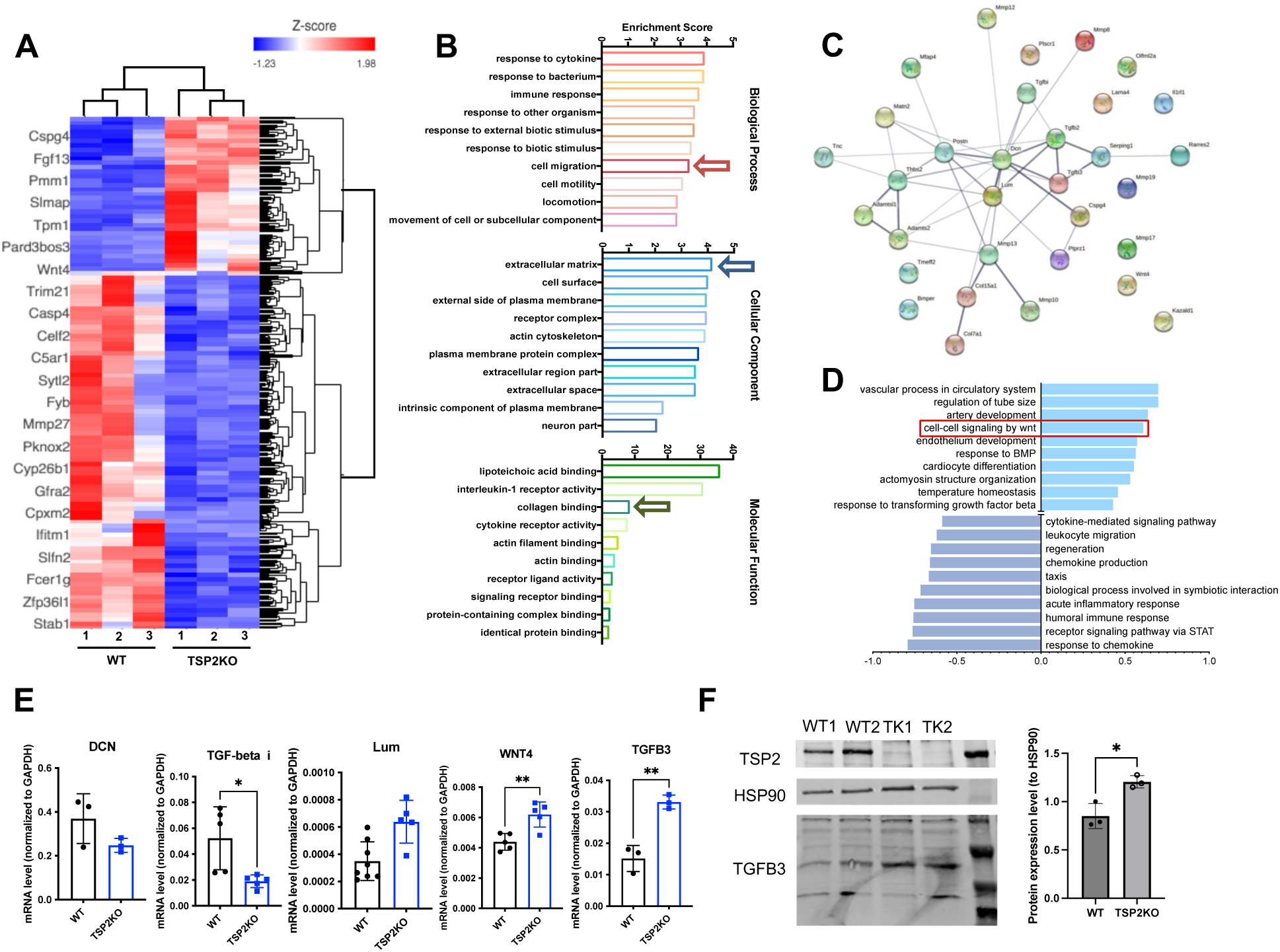
Bulk-RNA sequencing confirms perturbation in ECM proteins and suggests a positive regulation of Wnt pathway in KO fibroblasts. (A) Heatmap of genes that are differential expressed between two genotypes. (B) GO analysis. (C) Network map of ECM proteins identified. (D) GSEA analysis. (E) qPCR of genes that are shown up in multiple unbiased analysis in WT and KO primary fibroblasts. (F) Western blot of TGF-β3 in cell lysates isolated from WT and TSP2KO (TK) mouse dermal fibroblasts. (n = 3-8, unpaired T-test, two-tailed, *, p<0.05, **, p<0.001)

**Figure 5.**
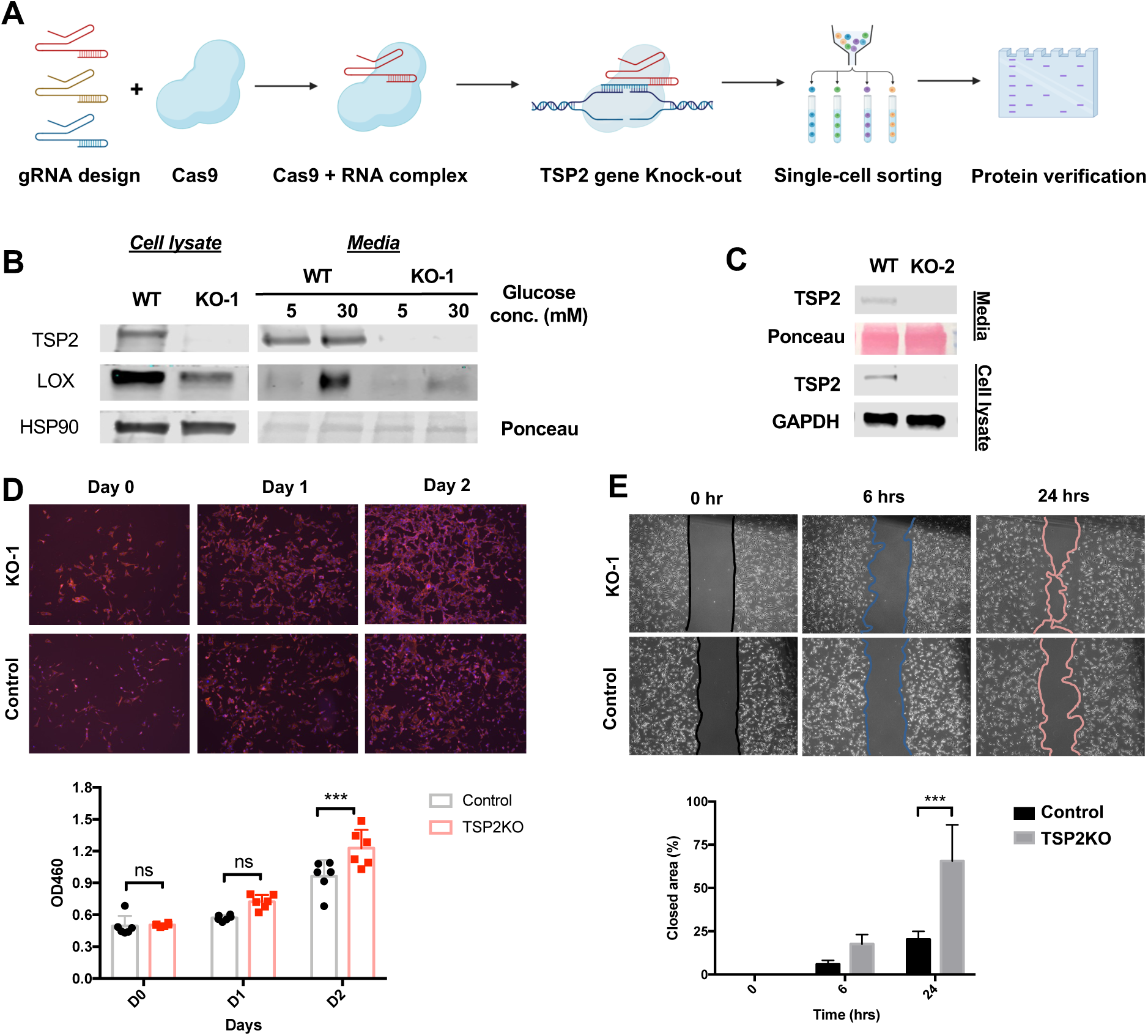
Depletion of TSP2 in NIH 3T3 cells using CRISPR/Cas9 creates a clean system to study function and pathway. (A) Schematic diagram of TSP2KO CRISPR NIH3T3 cell lines. Validation of (B) KO-1 and (C) KO-2 cell lines using western blot. (D) Proliferation and (E) scratch assay of KO and Control cells (n = 3-6, Unpaired T-test, two-tailed, ***, p<0.0001).

We then explored the impact of TSP2 deficiency on two critical cell functions in tissue repair— proliferation and migration, which were also highlighted as top regulated pathways from previous sequencing analysis. Cell proliferation, assessed using a CCK-8 assay, showed a higher cell count in the KO group compared to the control after 48 hours of culture (**Figure 5**). Similarly, scratch assay results revealed that TSP2 KO cells closed the wound area more efficiently than control cells 24 hours post-scratch (**Figure 5**). These in vitro analyses suggest that depletion of TSP2 improves fibroblast functions, further emphasizing the potential of TSP2 as a therapeutic target for impaired wound healing.

In summary, we successfully established two TSP2 KO CRISPR cell lines and observed enhanced proliferation and migration in the deficient cells. These findings align with previous *in vivo* observations and functional pathway analyses from RNA sequencing.

### TSP2 deficiency induces β-catenin activation in the setting of increased Wnt4/ TGF-β3

Pro-regenerative molecules and signaling were suggested to upregulated in TSP2 KO primary fibroblasts. Here, we reassured mRNA and protein levels of TGF-β3 and found it to be increased in TSP2KO NIH3T3 cells compared to controls (**Figure 6**). Moreover, WNT4, a WNT signaling pathway ligand, showed upregulation in CRISPR KO cells, which is consistent with observations in primary DFs (**Figure 4 & 6**). Generally, a β-catenin signaling cascade is initiated by the binding between WNT ligand and receptor with activation of Rac1. Previous research has also shown that TSP2 deficiency was associated with increased active Rac1 level. Therefore, we measured β-catenin in TSP2KO NIH3T3 cells to assess the activation state of WNT signaling. Immunofluorescence analysis showed an increased nuclear ratio of β-catenin in TSP2KO cells compared to WT and control 3T3 cells (**Figure 6**). Furthermore, western blot of Control and KO cell lysates confirmed increased level of total active β-catenin in the latter (**Figure 6**), suggesting a mechanism involving enhanced stabilization.

**Figure 6.**
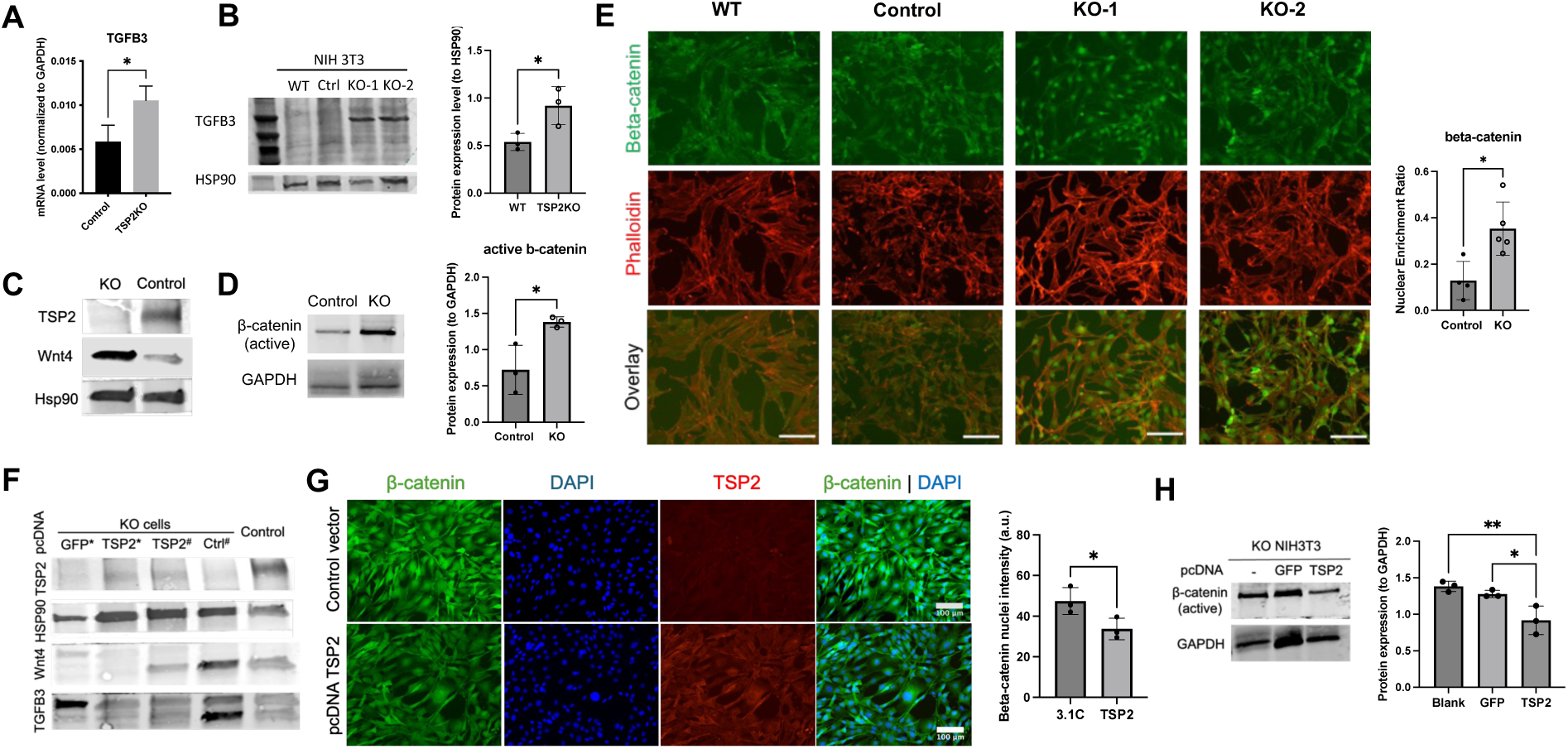
Depletion of TSP2 induces β-catenin activation with increased Wnt4/TGF-β3. (A) qPCR, (B) western blot and densitometrical analysis of TGF-β3. Western blot of (C) Wnt4 and (D) active β-catenin in Control and KO cells. (E) Immunofluorescence staining and quantification of nuclei enrichment ratio of β-catenin in WT, Control, KO-1 and KO-2 cells. (F) Western blot of Wnt4 and TGF-β3 in TSP2 rescued KO cells. (G) Immunofluorescence staining and nuclei intensity and (H) western blot of β-catenin in KO cells transfected with pcDNA3.1 c or pcDNA GFP and TSP2 (n=3-5, *, p<0.05, **, p<0.01).

Conversely, rescuing TSP2 expression in KO cells using pcDNA-TSP2 led to reduced Wnt4 and TGF-β3 protein levels compared to cells transfected with pcDNA-GFP (**Figure 6**). Furthermore, β-catenin immunostaining also showed lower nuclear intensity in KO cells transfected with pcDNA-TSP2 compared to those with pcDNA 3.1c.

In summary, TSP2 deficiency promotes β-catenin stabilization and nuclear translocation, accompanied by increased expression of Wnt4 and TGF-β3.

### Neurogenesis is enhanced in TSP2KO skin wounds with increased Sox10+ fibroblasts and TGF-β3 expression

Fibroblasts in TSP2KO skin are shown to shift towards a pro-regenerative phenotype including increased presence of Schwann multipotent Schwann cells, elevated levels of TGF-β3, and enhanced WNT signaling. Interestingly, a recent study indicates that β-catenin is required for the functional activation of Schwann cells following injury, leading to the release of TGF-β3, which promotes tissue repair [41]. Given the crucial role of dedifferentiated Schwann cells in peripheral nerve regeneration [41–45], we hypothesized that neurogenesis would be increased in TSP2KO skin wounds.

To test this hypothesis, we stained day 7 (D7) WT and TSP2KO skin wounds for Neurofilament H (NFH), a marker of neurons. Quantification of NFH+ areas within the total wound area revealed an increased formation of de novo nerve bundles in KO wounds compared to WT wounds (**Figure 7**). Additionally, the nerve bundles in KO wounds were significantly larger in diameter than those in WT wounds (**Figure 7**). Immunofluorescence staining for Sox10 in vivo demonstrated a higher number of Sox10+ cells in KO skin wounds (**Figure 7 and S6)**. Further analysis of TGF-β3+ areas confirmed increased production of TGF-β3 by KO fibroblasts (**Figure 7 and S6**).

**Figure 7.**
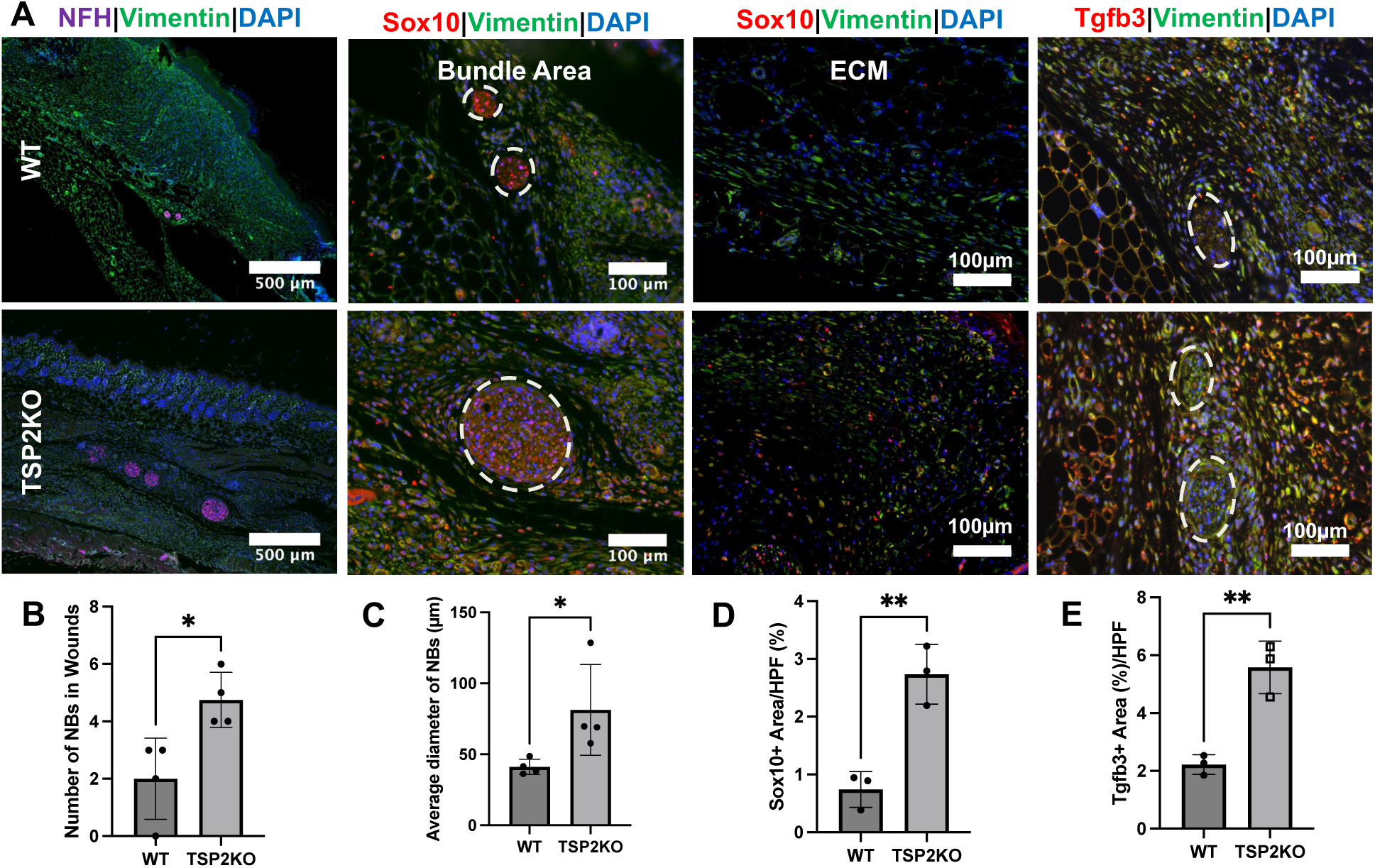
TSP2KO skin wounds demonstrate increased neurogenesis. (A) Immunofluorescence staining of NFH, Sox10, TGFb3 with Vimentin in D7 WT and TSP2KO mouse skin wounds. Quantification of (B) number of nerve bundles (NBs) in the center, (C) diameter of NBs (µm), (D) Sox10+ area/HPF (%), and (E) Tgfb3+ area/HPF (%) across the whole wounds (dashed circles indicate nerve bundles, n = 3-4, Unpaired T-test, two-tailed, *, p<0.05, **, p<0.01).

Taken together, these in vivo results support the hypothesis that depletion of TSP2 promotes neurogenesis in wounds. The increased production of TGF-β3, along with the higher presence of Sox10+ cells in KO wounds, not only validates in silico and in vitro observations, but also highlights a potential link between TSP2 deficiency and enhanced tissue regeneration. These findings suggest that TSP2KO skin environment is conducive to nerve repair, likely contributed by the shifted steady state of fibroblasts.

## Discussion

In this study, we explored the correlation between TSP2 and regeneration by focusing on fibroblasts, a major player in skin repair. Single-cell RNA sequencing enabled us to investigate fibroblast heterogeneity in greater detail. Despite differences between subtypes, all fibroblast populations shared common markers like Col1a1 and showed enrichment in pathways related to ECM proteins and cellular functions, consistent with current understanding of fibroblast biology. However, the overlapping expression of markers between fibroblast subtypes complicated cluster annotation in our study, particularly for the largest populations such as Dcn+, Rspo3+, and Dkk2+ fibroblasts. While these subtypes share common markers, certain genes were more highly expressed in specific clusters. For instance, Thy1, a key marker associated with fibrosis in skin [46], showed greater expression in Dcn+ fibroblasts. Thy1 is known to influence fibroblast lineage commitment between myofibroblasts and lipofibroblasts [47]. Similarly, En1, a marker more prevalent in Rspo3+ fibroblasts, has been reported to promote fibrosis during tissue repair. The reduction in both Dcn+ and Rspo3+ fibroblast populations in TSP2KO mice may contribute to the reduced scar formation in wounds and the reduced levels of Col1a1 protein in the KO ECM [25].

In addition to fibrotic markers, the traditional fibroblasts also express numerous ECM- and cell function-related proteins which are perturbed by TSP2 deficiency. For example, Ogn and Tnc, non-structural ECM proteins, are discovered to be the top down- and up-regulated genes in TSP2KO fibroblasts, respectively. Ogn, a proteoglycan, is known to modulate cell proliferation, migration, differentiation, and matrix assembly [48]. Knockdown of Ogn promotes smooth muscle cell proliferation and migration via the VEGF/VEGFR2 axis [49]. Conversely, overexpression of Tnc fosters fibroblast migration and differentiation, functions that are important for tissue repair [50]. We also observed changes in BMP pathway-related genes, such as the upregulation of Grem1, an antagonist of BMP signaling, and the downregulation of the transcription factor Hoxc8 in TSP2KO fibroblasts [34, 51, 52]. These alterations may contribute to the reduced BMP signaling identified in the cell-cell communication analysis, further highlighting the complex regulatory role of TSP2 in fibroblast behavior and ECM dynamics.

In contrast to the decrease of fibrosis-related fibroblast population, an increase of Sox10+ multipotent population is found in KO fibroblasts. While in vitro experiments have demonstrated that fibroblasts can be converted into Schwann cells with specific stimulus [53], the exact origin of these cells in the in vivo remains elusive. Nevertheless, KO ScP demonstrated elevated levels of PDGFb, Cdh2, and Jag1 signaling, which might contribute to their cell fate regulation via improved survival, enhanced migration, and inhibited differentiation. Specifically, PDGFb has been identified as a critical factor in the autocrine survival circuit of Schwann precursor cells [38, 54], and its enhanced expression in the KO Sox10+ population likely contributes to their wider distribution within the skin ECM. Moreover, Schwann precursor migration depends on interactions with both cells and the ECM, as they are directed by axonal guidance. A recent study identified N-cadherin (Cdh2) as a master regulator of collective Schwann cell migration through Slit2/3-mediated contact inhibition locomotion [55]. In our TSP2KO model, the increased expression of Cdh2 in ScPs, along with a more "loosened" ECM environment, likely facilitates their migration through the ECM. Additionally, elevated Jag1 is associated with activation of Notch signaling which has a dual role in Schwann cell biology: while it promotes proliferation in early stages, it acts as a negative regulator of myelination during later stages of differentiation [56, 57]. This suggests that Notch activation may contribute to the retention of a progenitor-like state in these cells. Interestingly, TSP2 has been reported to both potentiate and suppress Notch signaling through its interaction with Jag1 [58, 59], highlighting a potential regulatory mechanism that warrants further investigation. Despite the various hypotheses discussed above, the expansion of the Sox10+ population in TSP2KO fibroblasts is discovered and validated for the first time. These cells not only exhibit molecular similarities to multipotent Schwann progenitors but are also suggested to release pro-angiogenic and regenerative ligands. This profile may enhance the regenerative potential of the fibroblast population and foster an environment conducive to tissue repair.

Proliferation and migration, two cell functions that are critical for tissue repair, are enhanced in TSP2KO fibroblasts. The improvement in cell functions contribute to the acceleration of wound healing in TSP2KO mouse [21]. Under normal conditions, enhanced fibroblast proliferation and ECM deposition and contraction lead to scarring, which is not seen in the KO wounds. This suggests anti-scarring signaling pathways might also be activated with TSP2 depletion in addition to the deduction of fibrosis-related fibroblast population. One well-known anti-scarring molecule is TGF-β3 [60, 61], of which the expression level increases in KO fibroblasts and wounds. TGF-β3 exerts its anti-scarring effect via reducing collagen type 1 deposition and increasing MMP-9 production [61], which are also observed in TSP2KO wounds. Additionally, TGF-β3 mediates dermal fibroblast and keratinocytes migration to serve as a traffic control [62]. Interestingly, studies have found that exogenous WNT ligand treatment increased TGF-β3 expression and stimulation with TGF-β3 lead to Wnt4 upregulation [63, 64], which was seen in our TSP2-deficient cells.

Nevertheless, current knowledge is very limited on the interactions between TSP2 and the WNT pathway with only one study in cancer cells reporting that TSP2 activates WNT signaling [65]. In that study, TSP2 was shown to inhibit β-catenin degradation to induce activation. However, in our study, we observe opposite correlation between TSP2 and activation status of β-catenin. Specifically, we found that TSP2 deficiency led to enhanced stabilization and nuclear translocation of β-catenin, indicating activation of the pathway. The elevated level of WNT4, and increased active Rac1 suggested previously might work synergistically to induce this change [66] [67, 68]. Additionally, scRNA sequencing analysis suggested a decrease of WNT antagonist Sfrp2 in KO fibroblasts, which might contribute to β-catenin activation as well. As one of the initial molecular responses to injury, WNT activation plays a critical role in would healing and inhibited signaling has been reported in impaired tissue regeneration [69]. Strategies to re-activate β-catenin have led to improved cell functions and wound healing [70]. The enhanced cell proliferation, migration, and improved tissue regeneration observed in the TSP2KO environment are likely mediated, in part, by the activation of the WNT/β-catenin pathway, highlighting TSP2 as a promising therapeutic target for addressing impaired healing.

## Conclusion

We identified distinct molecular and functional heterogeneity among fibroblasts in both WT and TSP2KO skin, with an enrichment of pro-regenerative phenotypes in the latter. Notably, we observed an increased presence of Sox10+ populations, resembling Schwann precursor cells, and a reduction of fibrosis-related populations in TSP2KO fibroblasts. The expansion of Sox10+ populations was further confirmed through immunostaining of tissues and cells. Pro-survival and pro-migratory molecules, such as PDGFb and Jag1, expressed by these multipotent progenitors, may contribute to a conducive environment for tissue regeneration, including increased neurogenesis. Despite the cell heterogeneity observed, pathways related to cell function and ECM signaling were identified as the top regulated categories linked to TSP2 deletion, highlighting TSP2’s role as a mediator of cell and ECM interactions. Additionally, this study is the first to identify the positive regulation of pro-regenerative molecules and pathways, including TGF-β3 and WNT signaling, as downstream effects of TSP2 deficiency. The activation of β-catenin signaling likely contributes to the enhanced proliferation and migration seen in CRISPR-engineered TSP2KO fibroblasts. Together, changes in cell heterogeneity, improved cell functions, and activation of WNT4/β-catenin/TGF-β3 signaling likely underpin the enhanced tissue regeneration observed in TSP2KO wounds. These findings not only enhance our understanding towards fibroblasts in tissue repair but also highlight a potential therapeutic target for impaired regeneration.

## Data Availability

All data needed to evaluate the conclusions in the paper are included in this article and its supplementary information. Datasets are deposited onto NCBI GEO. Codes used for analysis are available upon request.

## Author Contribution Statement

Y.H. contributed to the design, conduction, data analysis, interpretation of all experiments. N.W. contributed to scRNA seq analysis. H.X. contributed to RNA seq data collection. J.T., J.Z., and J.L. contributed to CRISPR cell creation. D.G., H.C.H. contributed to flow cytometry. All authors provided feedback for the manuscript. S. R. contributed to data analysis and interpretation. T.R.K. contributed to project design and data interpretation. Y.H., T.R.K., and S. R. wrote the paper and all authors provided feedback.

### Acknowledgement

This project was supported by NIH grant DK132645 and National Institute of General Medical Sciences of the National Institutes of Health under Award Number 1S10OD030363-01A. Moreover, we want to thank Yale University Keck Microarray Shared Resource, Keck DNA Sequencing Facility and Yale Harvey Cushing/John Hay Whitney Medical Library for their assistance. We would also like to thank Dianqing (Dan) Wu Lab for sharing pcDNA 3.1c vector.

