## Supplementary Information for "Alteration of skin fibroblast steady state contributes to healing outcomes"

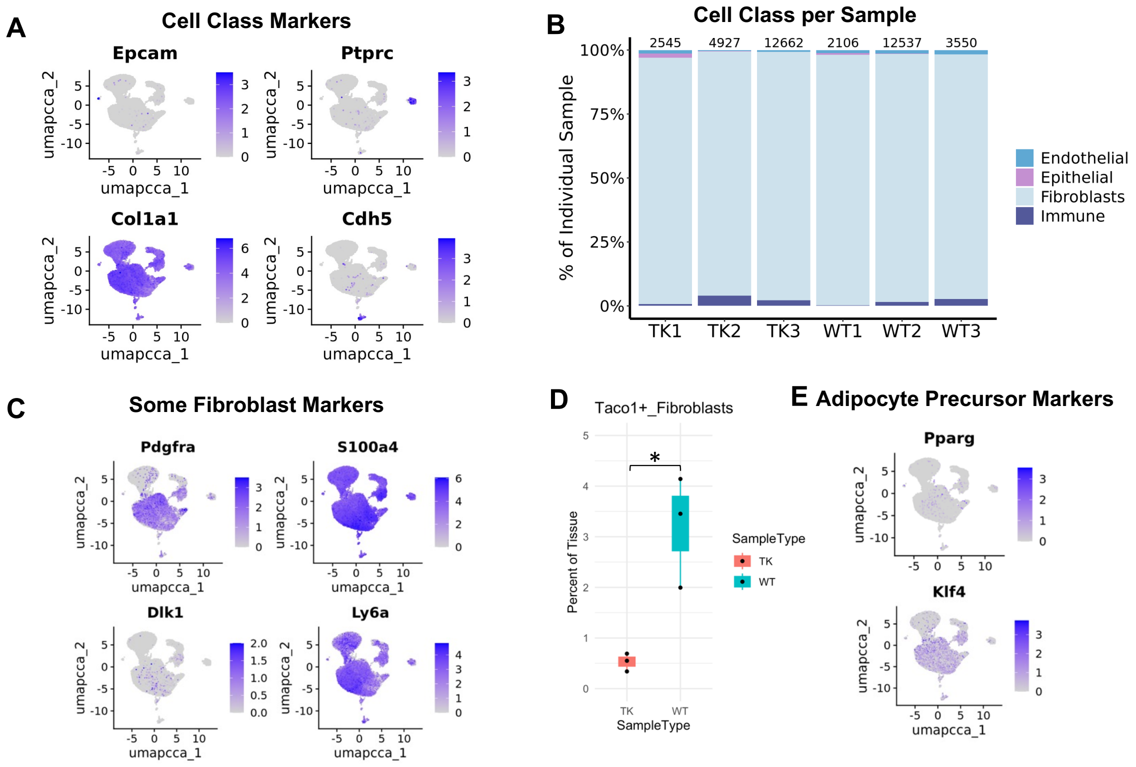


**Figure S1. Single cell transcriptomics reveals disctinct WT and TSP2KO dermal fibroblast heterogeneity.** (A) Feature plots of cell class markers: Epcam (Epithelial), Ptprc (Immune), Col1a1 (Fibro/Mesenchymal), Cdh5 (Endothelial); (B) Proportion of each class per sample; (C) Feature plots of some reported markers related to fibroblasts; (D) Comparison of Taco1+ fibroblasts percentage between WT and TSP2KO (TK) sample (Welch T-test, *, $p<0.05$); (E) Feature plots of Adipocyte Precursor related transcription factors.

In addition to the prominent differences observed in larger fibroblast populations, we also noted changes in smaller subsets that are of particular interest. Specifically, there was an increased presence of Ebf2+ cells and a diminished Taco1+ population in the TSP2KO group. Previous single-cell and spatial studies on early mouse skin development have identified the Ebf2+ population as muscle-related fibroblasts [54]. Given the role of Igfbp4 in regulating fat cell development [55], the enrichment of Igfbp4 in Ebf2+ fibroblasts may suggest a role for this population in energy-related lineage development. Conversely, the Taco1+ fibroblast population are enriched with Slit-Robo signaling molecules including Slit3 and Robo1, which may contribute to Robo1-related communication in WT. Additionally, interferon-related genes and immune responses were globally regulated in the bulk RNA-seq data, possibly due to the changes in IFN Stimulated fibroblasts.

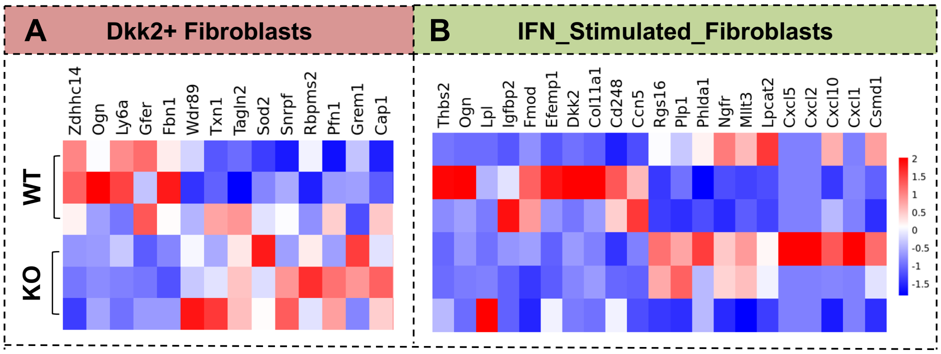


**Figure S2.** Heatmap of highly differential genes of (A) Dkk2+ (score ≥ 6, |power| > 1) and (B) IFN_Stimulated fibroblasts ((score ≥ 6, |power| > 10).

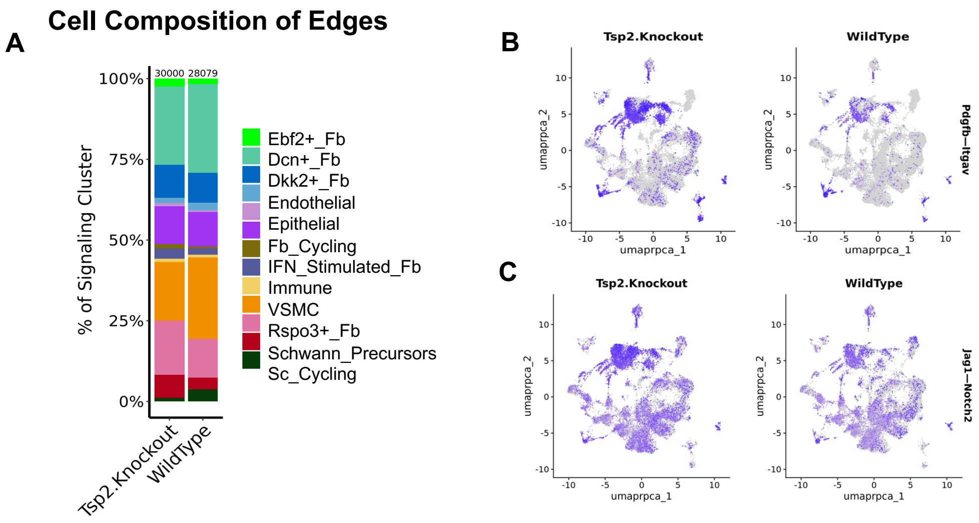


**Figure S3. NICHE analysis indicates an enhanced PDGFb and Notch signaling in TSP2KO fibroblast population.** (A) Cell proportion of NICHES signaling in WT and TSP2KO after downsampling; FeaturePlot of (B) Pdgfb – Itgav and (C) Jag1 – Notch signaling splited by genotype.

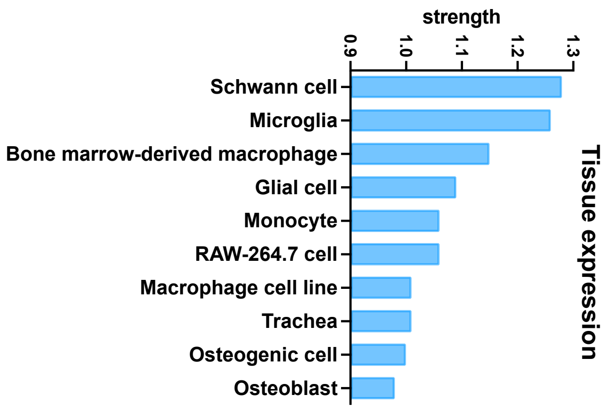


**Figure S4.** Tissue expression analysis on 462 genes that are differentially expressed between WT and TSP2KO fibroblasts.

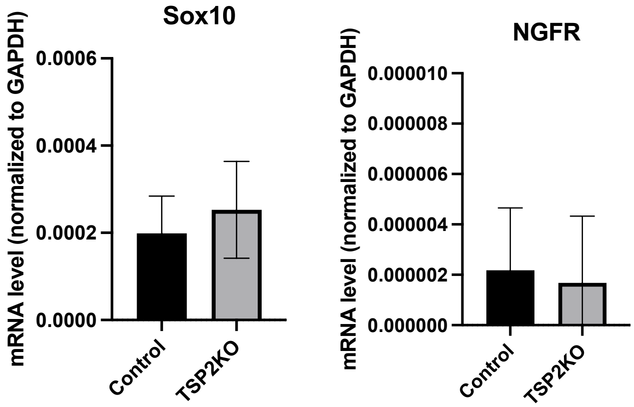


**Figure S5. CRISPR Knock out TSP2 in NIH 3T3s does not change cell state.** Sox10 and NGFR qPCR on Control and TSP2KO CRISPR cells. (n=3)

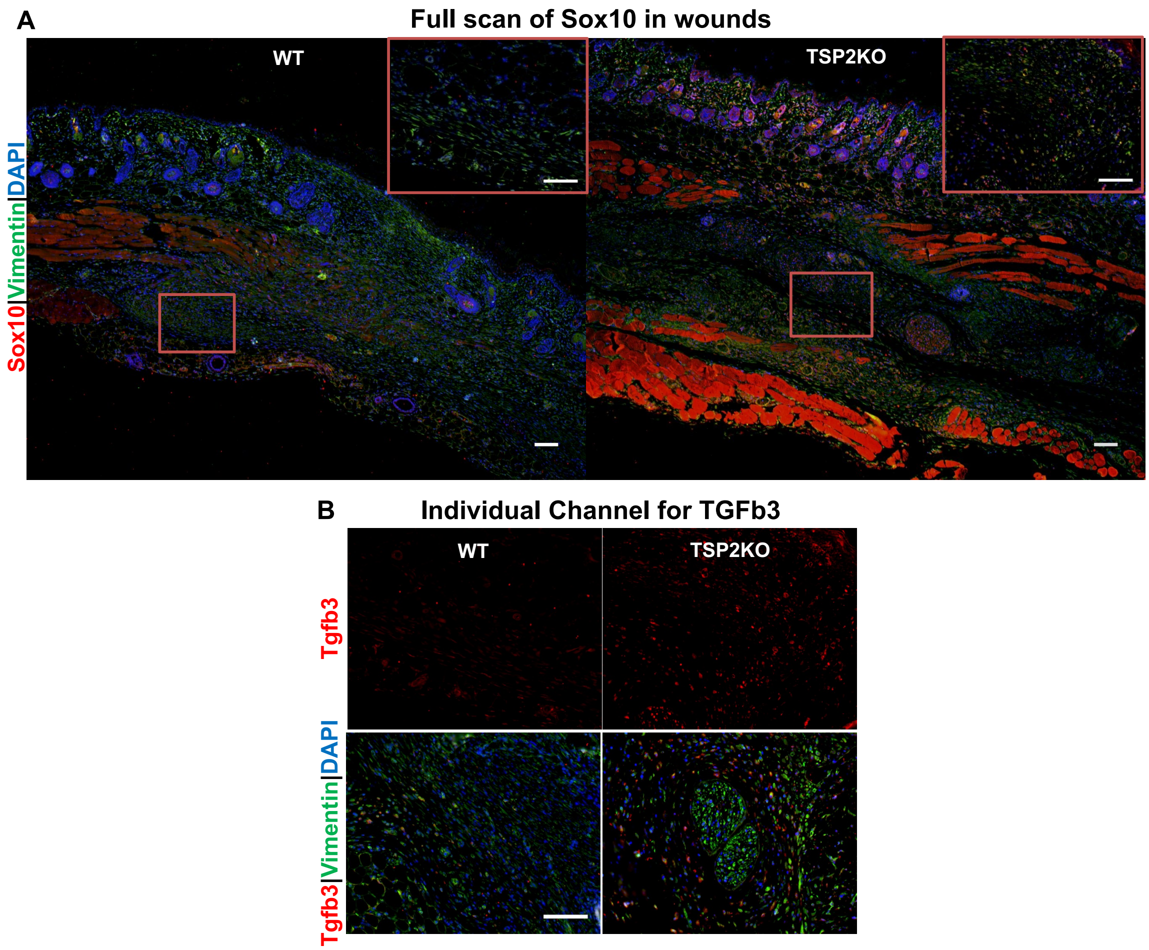


**Figure S6. TSP2KO skin wounds have increased Sox10 and TGfb3 expression.** (A) Representative images of the full scan of Sox10 staining in D7 WT and TSP2KO wounds. (B) Representative images of individual channel for TGFb3 staining shown in Figure 5 (scale bar = 100 μm).

**Table S1.** Sequences for Primer used

| Genes | Primer Sequence |
|---|---|
| TSP2 gRNA (KO-1) | 5’ - CACCG AAGCAGGACCGCAAGTCTCG - 3’<br>3’ - CTTCGTCCTGGCGTTCAGAGC CAAA - 5’ |
| TSP2 gRNA (KO-2) | 5’ - CACCG TCACATCACCACAGATCTCG - 3’<br>3’ - CAGTGTAGTGGTGTCTAGAGC CAAA - 5’ |
| DCN | F: ACTCTCCAGGAACTTCGTGTCC |

| | |
|---|---|
| | R: AGTCCCTGGAAGGCTCCGTTTT |
| TGFBI | F: AGCTGCTTATCCCAGATTCAGCCA<br>R: TATCGAGGCCAGCTTGTTTGAGGA |
| NGFR | F: GGAGAGAAACTGCACAGCGACA<br>R: CAGGCTACTGTAGAGGTTGCCA |
| Sox10 | F: AGATCCAGTTCCGTGTCAATAA<br>R: GCGAGAAGAAGGCTAGGTG |
| β-actin | F: CATTGCTGACAGGATGCAGAAGG<br>R: TGCTGGAAGGTGGACAGTGAGG |
| Lum | F: CCTGGAGGTCAATGAACTTG<br>R: CACTCATACATGTCAGGAGG |
| Wnt4 | F: ATCTCTTCAGCAGGTGTGGC<br>R: ACCCGCATGTGTGTCAAGAT |
| TGFB3 | F:AAGCAGCGCTACATAGGTGGCA<br>R:GGCTGAAAGGTGTGACATGGAC |
| GAPDH | F: TTCACCACCATGGAGAAGGC<br>R: GGCATGGACTGTGGTCATGA |

**Table S2.** Genes that are highly differential expressed between WT and KO Shwann Precursors (score ≥ 6, |power| >1, positive value indicates enrichment in KO)

| Genes | Power | Annotation |
|---|---|---|
| Pdgfb | 16.1270925 | Platelet-derived growth factor subunit B; Growth factor that plays an essential role in the regulation of embryonic development, cell proliferation, cell migration, survival and chemotaxis. |
| Mcam | 5.52074451 | Cell surface glycoprotein MUC18; Plays a role in cell adhesion, and in cohesion of the endothelial monolayer at intercellular junctions in vascular tissue. |
| Moxd1 | 3.25712345 | DBH-like monooxygenase protein 1; Belongs to the copper type II ascorbate-dependent monooxygenase family. |
| Jag1 | 3.21141252 | Protein jagged-1; Ligand for multiple Notch receptors and involved in the mediation of Notch signaling. |
| Plp1 | 2.42866167 | Myelin proteolipid protein; This is the major myelin protein from the central nervous system. |
| Cdh2 | 2.3213642 | Cadherin-2; Calcium-dependent cell adhesion protein; preferentially mediates homotypic cell-cell adhesion by dimerization with a CDH2 chain from another cell. |
| Col18a1 | 2.24088246 | Collagen alpha-1(XVIII) chain |
| Sdc4 | 2.06980442 | Syndecan-4; Cell surface proteoglycan that bears heparan sulfate. |
| Edil3 | 2.02770334 | EGF-like repeat and discoidin I-like domain-containing protein 3; Promotes adhesion of endothelial cells through interaction with the alpha-v/beta-3 integrin receptor. |
| Anxa1 | 1.60814944 | Annexin A1; Plays important roles in the innate immune response |
| Txn1 | 1.56385623 | Thioredoxin; Participates in various redox reactions |
| Ppfibp1 | 1.5084158 | Liprin-beta-1; May regulate the disassembly of focal adhesions |
| Rhob | 1.44223992 | Rho-related GTP-binding protein RhoB; Mediates apoptosis |
| Dpysl2 | 1.29249289 | Dihydropyrimidinase-related protein 2; Plays a role in neuronal development and polarity, as well as in axon growth and guidance, neuronal growth cone collapse and cell migration.. |
| Adam10 | 1.27147249 | Disintegrin and metalloproteinase domain-containing protein 10; Cleaves the membrane-bound precursor of TNF-alpha to its mature soluble form. |
| Smim15 | 1.20649584 | Small integral membrane protein 15. |
| Actn4 | 1.12520334 | Alpha-actinin-4; F-actin cross-linking protein which is thought to anchor actin to a variety of intracellular structures. |
| Tmsb4x | 1.1019539 | Hematopoietic system regulatory peptide; Plays an important role in the organization of the cytoskeleton. |
| Thbs2 | -10.4105566 | Thrombospondin-2 |
| Ly6a | -8.24492092 | Lymphocyte antigen 6A-2/6E-1; T-cell activation. |
| Rps2 | -1.15906657 | 40S ribosomal protein S2 |

**Table S3.** Genes that are highly differential expressed between WT and KO Dcn+ Fibroblasts (score ≥ 7, |power| >1, , positive value indicates enrichment in KO)

| Genes | Power | Annotation |
|---|---|---|
| Grem1 | 3.04308134 | Gremlin-1; Cytokine that may play an important role during carcinogenesis and metanephric kidney organogenesis. |
| Sod2 | 2.82525562 | Superoxide dismutase [Mn], mitochondrial; Belongs to the iron/manganese superoxide dismutase family. |
| H2-K1 | 1.60826344 | H-2 class I histocompatibility antigen, K-B alpha chain; Involved in the presentation of foreign antigens to the immune system. |
| Col18a1 | 1.84748236 | Collagen alpha-1(XVIII) chain |
| Tnc | 2.1471187 | Tenascin; Extracellular matrix protein implicated in guidance of migrating neurons as well as axons during development. |
| Postn | 1.55069083 | Periostin; Induces cell attachment and spreading and plays a role in cell adhesion. |
| Col5a3 | 2.42568937 | Collagen type V alpha 3 chain. |
| Cebpb | 1.57414331 | CCAAT/enhancer-binding protein beta; Important transcription factor regulating the expression of genes involved in immune and inflammatory responses. |
| Snrpf | 1.64584145 | Small nuclear ribonucleoprotein F; Plays role in pre-mRNA splicing |
| H2-D1 | 1.48560062 | H-2 class I histocompatibility antigen, D-B alpha chain |
| Thbs2 | -4.70344528 | Thrombospondin-2 |
| Ogn | -3.93797233 | Mimecan; Induces bone formation in conjunction with TGF-beta-1 or TGF- beta-2; Belongs to the small leucine-rich proteoglycan (SLRP) family. SLRP class III subfamily. |
| Igfbp2 | -2.86019706 | Insulin-like growth factor-binding protein 2; Inhibits IGF-mediated growth and developmental |
| Dkk2 | -2.35942511 | Dickkopf-related protein 2; Antagonizes canonical Wnt signaling |
| Col4a5 | -3.06569439 | Collagen, type IV, alpha 5. |
| Col8a1 | -2.06376439 | Collagen alpha-1(VIII) chain |
| Zdhhc14 | -2.21030915 | Probable palmitoyltransferase ZDHHC14; Belongs to the DHHC palmitoyltransferase family. ERF2/ZDHHC9 subfamily. |
| Efemp1 | -1.97269991 | EGF-containing fibulin-like extracellular matrix protein 1; Binds EGFR |
| Zfp52 | -2.72141969 | Zinc finger protein 52. |
| St3gal5 | -1.75786222 | Lactosylceramide alpha-2,3-sialyltransferase; Catalyzes the formation of ganglioside GM3 |
| Gfer | -1.7770521 | FAD-linked sulfhydryl oxidase ALR; FAD-dependent sulfhydryl |
| Col11a1 | -1.63699153 | Collagen alpha-1(XI) chain |
| S100a1 | -1.90185829 | Protein S100-A1; Probably acts as a Ca(2+) signal transducer |
| Hoxc8 | -1.41981781 | Homeobox protein Hox-C8; Sequence-specific transcription factor |

| | | |
|---|---|---|
| Prelp | -1.35107422 | Prolargin |
| Gria3 | -1.12937304 | Glutamate receptor 3; Receptor for glutamate |
| Id4 | -1.51338676 | DNA-binding protein inhibitor ID-4. |
| Sorbs1 | -1.60946 | Sorbin and SH3 domain-containing protein 1. |
| Fgf10 | -1.31480532 | Fibroblast growth factor 10. |

**Table S4.** Genes that are highly differential expressed between WT and KO Rspo3+ Fibroblasts (score ≥ 7, |power| >1, positive value indicates enrichment in KO)

| Genes | Power | Annotation |
|---|---|---|
| Cap1 | 2.34312314 | Adenylyl cyclase-associated protein 1; Directly regulates filament dynamics. |
| Cebpb | 1.80219021 | CCAAT/enhancer-binding protein beta; transcription factor |
| Snrpf | 1.65656915 | Small nuclear ribonucleoprotein F; Plays role in pre-mRNA splicing |
| Pfn1 | 1.35448171 | Profilin-1; Binds to actin and affects the structure of the cytoskeleton. |
| Tnc | 1.8015762 | Tenascin. |
| Sod2 | 1.85721203 | Superoxide dismutase [Mn], mitochondrial |
| Eef1e1 | 1.88792561 | Eukaryotic translation elongation factor 1 epsilon-1; Positive modulator of ATM response to DNA damage. |
| Cks2 | 1.66616649 | Cyclin-dependent kinases regulatory subunit 2 |
| Plk2 | 1.78237292 | Serine/threonine-protein kinase PLK2. |
| Tgif1 | 1.67093347 | Homeobox protein TGIF1; Binds to a retinoid X receptor (RXR) |
| Txn1 | 1.25953222 | Thioredoxin; Participates in various redox reactions |
| Snhg6 | 1.25169621 | Small nucleolar RNA host gene 6, ncRNA |
| Tfrc | 1.30417984 | Transferrin receptor protein 1 |
| Trp53 | 1.54240631 | Cellular tumor antigen p53; Acts as a tumor suppressor |
| Sfrp1 | -7.82621157 | Secreted frizzled-related protein 1; Soluble frizzled-related proteins (sFRPS) function as modulators of Wnt signaling through direct interaction with Wnts. |
| Thbs2 | -4.60638529 | Thrombospondin-2 |
| Ogn | -6.58432231 | Mimecan; Induces bone formation in conjunction with TGF-beta |
| Tshz2 | -3.76243072 | Teashirt homolog 2; Probable transcriptional repressor |
| Etl4 | -3.19852729 | Sickle tail protein |
| Col11a1 | -2.15924824 | Collagen alpha-1(XI) chain; May play an important role in fibrillogenesis |
| Slit2 | -1.40500808 | Slit homolog 2 protein C-product |
| Itpr1 | -1.61404058 | Inositol 1,4,5-trisphosphate receptor type 1 |
| Hoxc8 | -1.5319403 | Homeobox protein Hox-C8; Sequence-specific transcription factor |
| Ebf1 | -1.09487703 | Transcription factor COE1; Transcriptional activator |
| St3gal5 | -1.32147436 | Lactosylceramide alpha-2,3-sialyltransferase |
| Slc9a3r2 | -1.48716904 | Na(+)/H(+) exchange regulatory cofactor NHE-RF2 |

**Table S5.** Genes that are highly differential expressed between WT and KO Dkk2+ Fibroblasts (score ≥ 6, |power| >1, positive value indicates enrichment in KO)

| **Genes** | **Power** | **Annotation** |
|---|---|---|
| Cap1 | 2.16555204 | Adenylyl cyclase-associated protein 1; Directly regulates filament dynamics. |
| Grem1 | 2.36522544 | Gremlin-1; as BMP antagonist |
| Pfn1 | 1.60881226 | Profilin-1; Binds to actin and affects the structure of the cytoskeleton. |
| Rbpms2 | 1.8818222 | RNA-binding protein with multiple splicing 2.Mediates an increase of NOG mRNA levels, thereby contributing to the negative regulation of BMP signaling pathway |
| Snrpf | 1.7465596 | Small nuclear ribonucleoprotein F; Plays role in pre-mRNA splicing |
| Sod2 | 1.87444531 | Superoxide dismutase [Mn], mitochondrial. |
| Tagln2 | 1.12113149 | Transgelin-2; Belongs to the calponin family. |
| Txn1 | 1.1035159 | Thioredoxin; Participates in various redox reactions |
| Wdr89 | 1.05217188 | WD repeat-containing protein 89. |
| Ogn | -8.38311408 | Mimecan; Induces bone formation in conjunction with TGF-beta |
| Zdhhc14 | -2.27194489 | Probable palmitoyltransferase ZDHHC14; Belongs to the DHHC palmitoyltransferase family. |
| Ly6a | -2.68180966 | Lymphocyte antigen 6A-2/6E-1; T-cell activation. |
| Fbn1 | -1.34326028 | Fibrillin-1; [Fibrillin-1]: Structural component of the 10-12 nm diameter microfibrils of the extracellular matrix |
| Gfer | -1.15290088 | FAD-linked sulfhydryl oxidase ALR. |

**Table S6.** Genes that are highly differential expressed between WT and KO IFN Stimulated Fibroblasts (score ≥ 6, |power| >10, positive value indicates enrichment in KO)

| **Genes** | **Power** | **Annotation** |
|---|---|---|
| Cxcl1 | 3024.20748 | Growth-regulated alpha protein; Has chemotactic activity for neutrophils. |
| Cxcl10 | 42.3476581 | C-X-C motif chemokine 10; Pro-inflammatory cytokine |
| Phlda1 | 29.3359672 | Pleckstrin homology-like domain family A member 1; Seems to be involved in regulation of apoptosis. |
| Lpcat2 | 13.7535215 | Lysophosphatidylcholine acyltransferase 2; Possesses both acyltransferase and acetyltransferase activities |
| Plp1 | 12.7885131 | Myelin proteolipid protein; |
| Mllt3 | 15.2097363 | Protein AF-9; Chromatin reader component of the super elongation complex (SEC). |
| Ngfr | 10.0988291 | Tumor necrosis factor receptor superfamily member 16; Low affinity neurotrophin receptor which can bind to mature NGF, BDNF, NTF3, and NTF4. |
| Rgs16 | 14.691323 | Regulator of G-protein signaling 16; Regulates G protein-coupled receptor signaling cascades. |

| | | |
|---|---|---|
| Csmd1 | 10.9171791 | CUB and sushi domain-containing protein 1; Belongs to the CSMD family. |
| Dkk2 | -34.0617306 | Dickkopf-related protein 2; Antagonizes canonical Wnt signaling. |
| Ogn | -31.0339248 | Mimecan; Induces bone formation in conjunction with TGF-beta |
| Igfbp2 | -19.4448737 | Insulin-like growth factor-binding protein 2. |
| Efemp1 | -19.3589908 | EGF-containing fibulin-like extracellular matrix protein 1 |
| Ccn5 | -18.5147083 | CCN family member 5; May play an important role in modulating bone turnover. |
| Lpl | -12.8474573 | Lipoprotein lipase; Key enzyme in triglyceride metabolism. |
| Col11a1 | -11.9664641 | Collagen alpha-1(XI) chain. |
| Fmod | -13.5473518 | Fibromodulin; Affects the rate of fibrils formation. |
| Cd248 | -13.6844605 | Endosialin; May play a role in angiogenesis or vascular function. |
| Thbs2 | -13.0784781 | Thrombospondin-2 |
| Srpx | -12.6538612 | Sushi Repeat Containing Protein X-Linked |
